# Interaction of levothyroxine with bovine serum albumin: a spectroscopic assay

**DOI:** 10.1101/2020.04.29.068510

**Authors:** Nicoleta Sandu, Claudia G. Chilom, Melinda David, Monica Florescu

## Abstract

Bovine serum albumin (BSA) acts as a carrier for many endogenous and exogenous compounds, such as thyroid hormones or corresponding drugs. Binding of the hydrophilic compound levothyroxine (LT4) to BSA can significantly alter the pharmacological properties of the compound. Therefore, studying its interaction with BSA could be a difficult issue. In this work, the binding mechanism and affinity of the interaction between LT4 and BSA were investigated, both in solution using UV-Vis, Fourier-transform infrared spectroscopy (FT-IR), fluorescence and fluorescence resonance energy transfer (FRET), as well as by Surface Plasmon Resonance (SPR) with BSA confined to a gold-coated chips, as far as we know for the first time used to study the interactions between LT4 and proteins. Quenching of BSA fluorescence by LT4 combined with UV-Vis spectroscopy shows a ground-state complex formation that may be accompanied by a nonradiative energy transfer process. FT-IR revealed the changes induced by LT4 in the secondary structure of BSA molecules, due to the partial unfolding of BSA native structure upon LT4 binding. Scatchard approach allowed the determination of the binding constant and the thermodynamic parameters, which correspond to an enthalpic process, driven mainly by hydrogen bonds and van der Waals forces. Using SPR, the adsorbed amount of biomolecules was calculated and the binding affinity of LT4 with confined-BSA was characterized using the Hill-Langmuir equation, indicating that the BSA immobilization plays an important role in LT4 binding. As preliminary results, both fluorescence quenching and SPR can be used as a stepping stone for the development of a spectroscopic biosensor for LT4 detection, with a limit of detection as low as 0.23 × 10^−6^ M.

## 1. Introduction

The body’s natural thyroid hormones are thyroxine (T4) and triiodothyronine (T3). They are produced by the thyroid, under the control of the pituitary thyroid stimulating hormone (TSH). Free thyroid hormones (that are metabolically active at the tissue level) are transported through the blood by serum proteins: thyroxine-binding globulin (TBG), transthyretin (TTR), and albumin (1, 2). Then, thyroid hormones bind to protein receptors located on the surface of cell membranes (3) or to membranes of the mitochondria and nucleus (4). As a result, some specific biochemical processes are activated that produce important metabolic effects at the cellular level.

Most cellular and tissue functions are influenced by thyroid hormones. Thus, thyroid hormones influence the production and breakdown of steroid hormones (5), the metabolism of carbohydrates (6, 7), fats (8), and proteins (9). Also, thyroid hormones specifically affect the development of the brain (10), heart rate (11, 12), pulmonary blood irrigations (13), but also some processes of the immune system (14). Growth, development and reproduction of the body are influenced by thyroid hormones (15, 16).

The reversible transformation of the two thyroid hormones (i.e., T3 and T4) results in a very fine balance, which contributes to the proper functioning of the cellular processes or can lead to developmental problems and even cancer. This balance is influenced by many factors, including living in areas with low/high iodine content (17). Thyroid disorders are associated with the risk for breast, prostate, lung, ovary, gastric, pancreas, thyroid, and other cancer (18), and for some of them, the best therapy is the hormone replacement. Levothyroxine (LT4) is the pharmaceutical name of the manufactured version of T4. Herewith, the monitoring of LT4 levels in the human organism while under treatment is of great importance.

Albumins are the major soluble proteins in the blood which act as a deposit and also a carrier for many endogenous and exogenous compounds, such as drugs, including LT4. In drug-protein interaction studies, BSA is the most widely used protein due to its structural homology to human albumin (HSA), it is cheaper, and much more stable. BSA structure and function depend on a number of factors, such as temperature and pH (19) or association of ligands such as drugs (20), nanoparticles (21), dyes (22) or antioxidants such as vitamins and flavonoids (23–25). Binding of drugs to albumin causes a decrease in the concentration of the free form of the drug but also causes an increase in its half-life in plasma. Binding of the drug, such as LT4, to BSA can significantly alter the pharmacological properties of the compound. As a hydrophilic compound, LT4 has a low binding of serum proteins and, depending on the methods used for testing, its binding varies. Therefore, studying its interaction with BSA could be a difficult issue. Therefore, understanding the pattern and mechanism of LT4-BSA interaction is very important for clinical care and evaluation of the optimal dose and clearance of the drug in the body.

The aim of this paper is to investigate the binding mechanism and affinity between LT4 and BSA in two situations, with BSA in free and surface-confined form. The methods chosen for this investigation were UV-Vis and FT-IR spectroscopy, fluorimetry, as well as SPR. SPR enables the study of label-free biomolecular interaction processes, highlighting small molecules binding by monitoring the small changes in the refractive index in the vicinity of a metal surface. While standard spectroscopic methods use the free form of BSA molecules, in SPR these molecules must be immobilized on a solid substrate (gold surface chip). To our knowledge, there is no other study in the literature that addresses the interaction between LT4 and BSA using SPR. The following parameters were determined: the strength of the interaction (monitored by the binding and affinity constants), the stoichiometry (the number of protein binding sites for the ligand) and the thermodynamic parameters. By binding to transporter proteins (including albumin), thyroid hormones become inactive. Thus, understanding the binding mechanism, which depends on changing the concentration of transporter proteins and environmental conditions, is important for the optimal circulation of hormones through plasma.

## 2. Materials and methods

### 2.1 Materials

Bovine serum albumin (purity over 98 %) was purchased from SIGMA, and its concentration was measured with a Perkin Elmer Lambda 2S spectrophotometer, using the standard molar absorption coefficient for Trp and Tyr at 280 nm (*ε* = 44,000 M^−1^ cm^−1^). Levothyroxine sodium pentahydrate, LT4 (888.93 g mol^−1^) solubilized in dimethyl sulfoxide (DMSO) to a stock solution of 2.8 mM was purchased from SIGMA. Glutaraldehyde 25 % (GA), ethanolamine 1 % (ETA), N-hydroxysuccinimide (NHS), N-ethyl-N3-dimethylaminopropyl carbodiimide (EDC), Cystamine (Cys) and N-(2-Hydroxyethyl)piperazine-N′-(2-ethanesulfonic acid) (HEPES ≥ 99.5 %) were also purchased from SIGMA. All samples were prepared in 0.1 M HEPES buffer, adjusted to pH 7.4 with a saturated NaOH solution. Millipore Milli-Qnanopure water (resistivity ≥ 18 MΩ cm) was used for the preparation of all solutions.

### 2.2 Methods

#### UV-Vis spectroscopy

UV spectra were recorded with a Perkin-Elmer spectrophotometer, in a 1 cm x 1 cm quartz cuvette, with 500 nm/min. All measurements were carried out at room temperature 24 °C ± 1 °C.

#### Fourier-transform infrared spectroscopy (FT-IR)

FT-IR spectra were recorded with a Perkin Elmer spectrometer, over the spectral range, 400 - 4,000 cm^−1^, with 4 cm^−1^ spectral resolution and 64 scans per sample. All samples were deposited on the silica plates and dried in a nitrogen atmosphere to evaporate the aqueous solvent. All measurements were carried out at room temperature 24 °C ± 1 °C.

#### Fluorescence spectroscopy

Fluorescence emission spectra of the BSA and BSA-LT4 samples were recorded using Perkin Elmer MS 55 spectrofluorometer, in the spectral range, 300 - 500 nm, with 500 nm/min speed, at an excitation wavelength of 295 nm. The slits for the excitation and emission monochromators were 4.0 nm and 4.5 nm, respectively. The BSA fluorescence was quenched by successive additions of LT4 (0 – 10) μM in the protein solution (3 μM), at 25 °C and 35 °C, with 5 spectral accumulations. The solutions were mixed and kept 3 min before measurements. All measurements were carried out at room temperature 24 °C ± 1 °C.

#### Fluorescence resonance energy transfer (FRET)

The distance between BSA and LT4 molecules, was also investigated by FRET. The efficiency of energy transfer (*E*) was calculated using the eq. (1) (26):

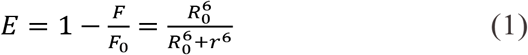

where *F_0_* and *F* are the fluorescence intensity of the donor (the protein) in the absence and respectively, presence of the acceptor (the ligand), *r* is the distance between the bound ligand on protein and the Trp residue, and *R_0_* (in Ǻ) is the Förster critical distance at which 50 % of the excitation energy is transferred to the acceptor. *R_0_* can be calculated by the following eq. (26):

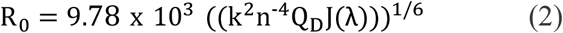

where *k* is describes the relative orientation of the transition dipoles of the donor and acceptor, *n* is the refractive index of medium, *Q_D_* is the quantum yield of donor in the absence of acceptor, and *J* is the overlap integral of the fluorescence emission spectrum of the donor with the absorption spectrum of the acceptor. *J* can be calculated according to the eq. (3):

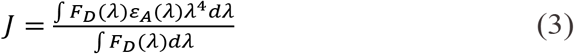

where *F_D_* (*λ*) is the corrected fluorescence intensity of the donor at wavelength *λ* with the total intensity (area under the curve) normalized to unity, *ɛ_A_* is the molar absorption coefficient (in M^−1^ cm^−1^) of the acceptor at the wavelength *λ.* All measurements were carried out at room temperature 24 °C ± 1 °C.

#### Surface Plasmon Resonance

A SPR analyzer system, MP-SPR NaviTM 200 OTSO, and planar gold SPR discs from BioNavis LTD, Finland, were used to evaluate the interactions between confined BSA and LT4, with the software packages for data acquisition and analysis SPR Navi Control and SPR Navi Data Viewer. The SPR analyzer is based on the Kretschmann configuration in which a polarized, collimated light beam undergoes total internal reflection at a glass/thin-metal-film/dielectric interface. It is equipped with 2 lasers with wavelength of 670 nm. Two types of chips were used and for both the surface is formed by a glass chip coated with a 50 nm thin gold layer which produces the SPR effect. One was used as such, and the other was functionalized with amino moieties (−NH_2_). The working temperature was 25 °C. All measurements were performed in a cell with two channels (1 μL per channel), in a continuous flow of 0.1 M HEPES, pH 7.4, with a flow rate of 30 μL/min throughout the whole experiment.

## 3. Results and discussions

Levothyroxine (LT4), the synthetic analogue of thyroxine T4 hormone, is used as the initial treatment for thyroid disease, especially hypothyroidism. The chemical structure of LT4 is shown in Scheme 1. The −NH_2_ group is the one that will be involved in a possible interaction with proteins, as we will discuss later.

**SCHEME 1.**
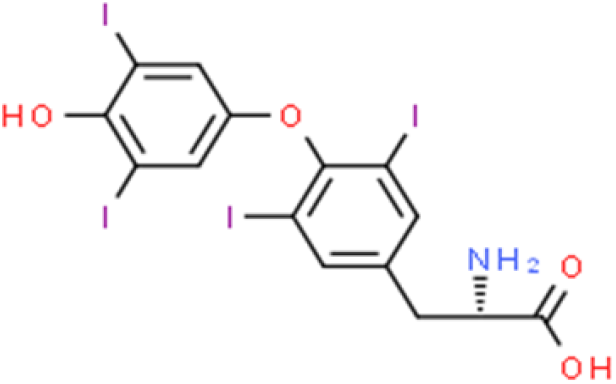
The chemical structure of levothyroxine (LT4) (27).

The stability and activity of biological molecules are influenced by factors as pH (19), temperature (28) or ligand interactions (22). The pH values of LT4 in saturated water solution are in the range of 8.35 to 9.35 (29). In order to determine the optimum pH value of LT4 diluted in HEPES buffer, the UV absorption spectrum of 20 μM LT4 was recorded. The maximum of the absorbance was found at 238 nm and a second maximum was obtained around 320 nm (Fig. 1A).

**FIGURE 1.**
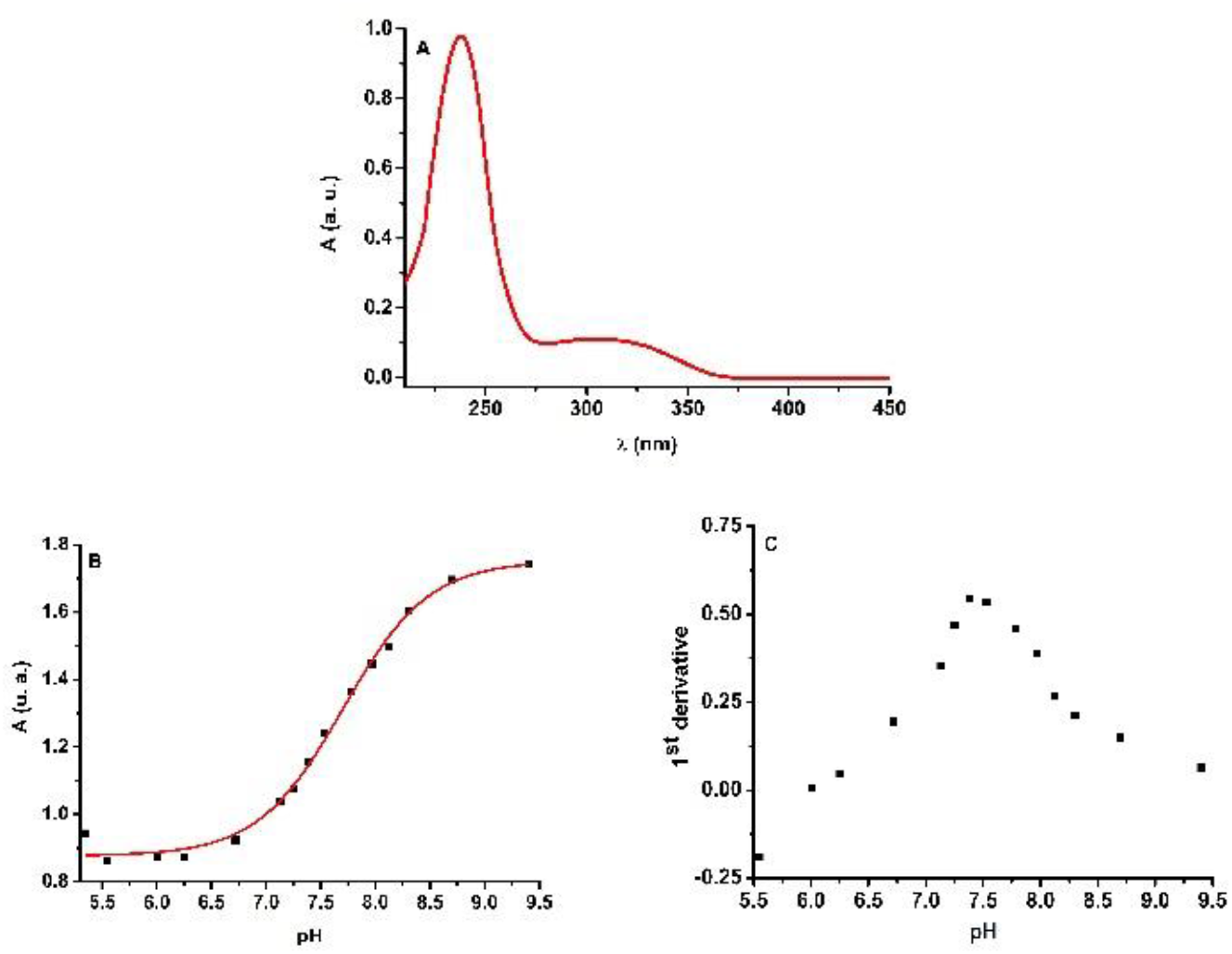
(A) The UV spectrum of LT4 diluted in HEPES buffer solution, pH 7.4. (B) The absorbance of LT4 at pH in the range of 5.35 and 9.40, measured at 240 nm. (C) The representation of the 1^st^ derivative of the absorbance.

### 3.1 Determination of LT4 optimal pH

There is no clear information in the literature on the optimal pH value at which LT4 can interact with proteins. For this reason, this study, the absorbance of LT4 at 238 nm was recorded at different pH values (5.35 - 9.35) (Fig. 1B) and the 1^st^ derivative of this representation (Fig 1C) was done. In this way, we found that the optimal pH value of LT4 in HEPES buffer is around 7.40. In order to work close to the physiological conditions, in the following spectroscopic experiments, LT4 was studied at pH 7.40.

### 3.2 The Binding of LT4 to BSA as measured by FT-IR spectroscopy

FT-IR spectroscopy is a useful tool to screen the binding interactions. The binding of LT4 to BSA was investigated on the basis of the involvement of the amino group in LT4 molecule (Fig 2A) and the amide I (C=O stretching) and amide II (C-N stretching coupled to NH bending) bands in BSA (30). The first signal corresponding to NH_2_ stretching in LT4 molecule is shifted from 2,954 cm^−1^ to 2,920 cm^−1^ in the BSA-LT4 complex, and the second one, from 2,836 cm^−1^ is completely attenuated in the BSA-LT4 complex. This is an indication that LT4 interact with BSA molecule *via* −NH_2_ group.

**FIGURE 2.**
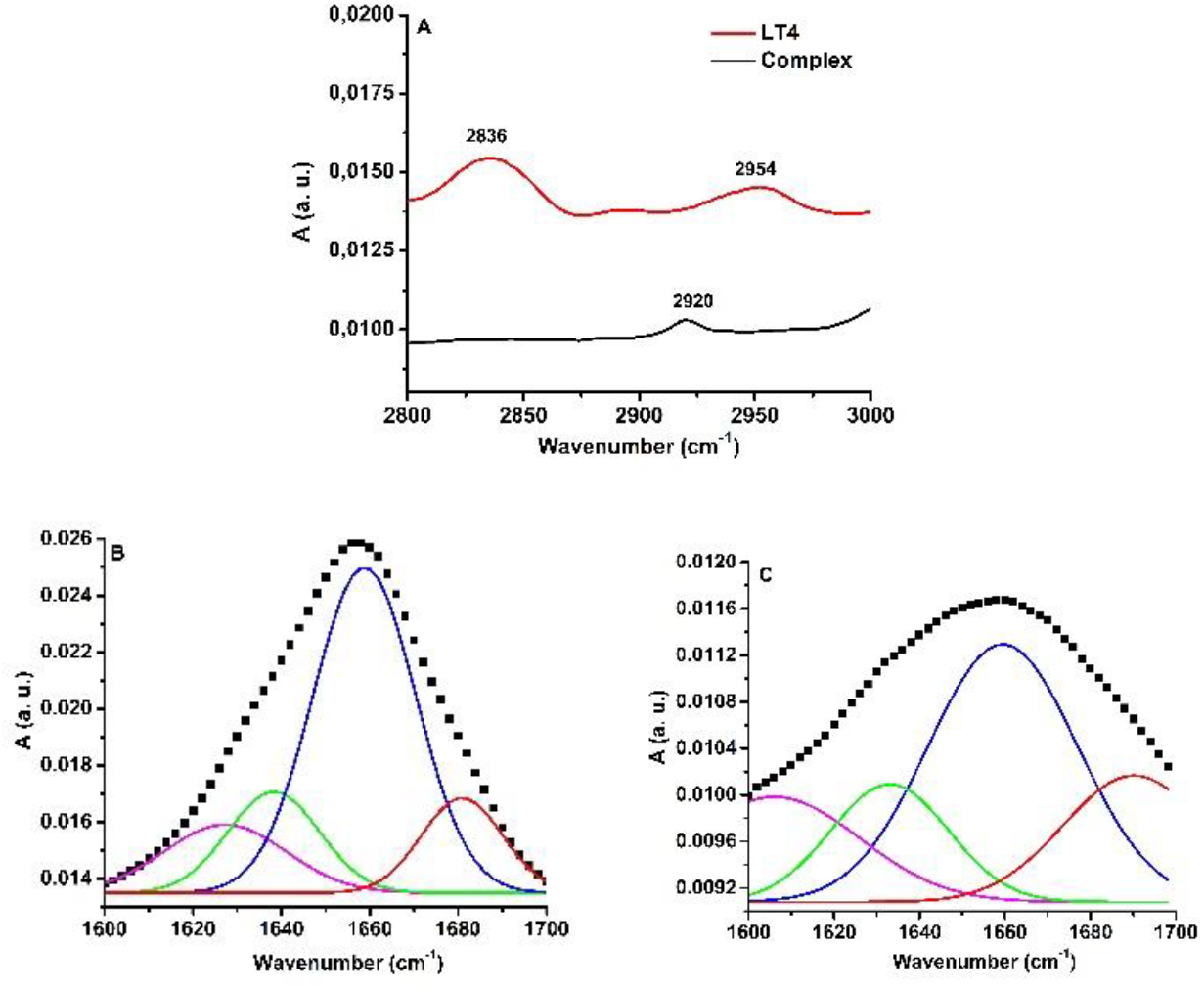
(A) FT-IR curves of LT4 and BSA-LT4 between the 3,000 – 2,800 cm^−1^, corresponding to the NH2 stretching in LT4 is shifted from 2,954 cm-1 to 2,920 cm-^1^ in the BSA-LT4 complex; the peak from 2836 cm^−1^ in LT4 is completely attenuated in the BSA-LT4 complex; curve-fitted amide I band between the 1,700 - 1,600 cm^−1^ for: (B) free BSA and (C) BSA-LT4. The amide I band is represented in black and the curves for the secondary structure elements are: blue for α helix, magenta for β turn in, red for β sheet and green for random coil.

The conformational changes after BSA-LT4 interactions were evaluated in the spectral range of 1,700 - 1,600 cm^−1^ (Fig 2B, C). A spectral shift of the amide I band from 1,656 cm^−1^ (free BSA) to 1,660 cm^−1^ for BSA-LT4 complex was observed. The spectral deconvolution shows that LT4 provokes changes in the BSA secondary structure elements. These values are presented in Table 1. The α helix content decreased from 56.94 % to 43.64 %, while the β sheet content increased from 14.18 % to 20.36 %. Also, the β turn content increased from 15.78 % to 20.35 %.

**TABLE 1.**
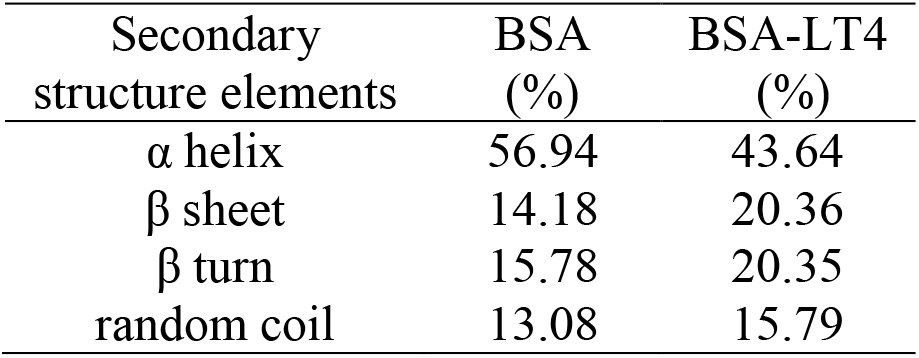
The percentages of secondary structure elements of BSA and BSA-LT4 complex for Amide I band.

### 3.3 The quenching of the intrinsic fluorescence of BSA by LT4

Albumin is one of the transport proteins for LT4 (31). For this reason, the understanding of the binding mechanism of LT4 to the albumin site is a starting point in the use of LT4 in living cells. The intrinsic fluorescence of BSA was used to investigate the binding mechanism of LT4 to BSA. The fluorescence emission of BSA (due to the Trp residues) was decreased by the gradual addition of LT4 in the protein solution. This behaviour is shown in Fig. 3A for the binding of LT4 to BSA monitored at 25 °C. A similar result was obtained at 35 °C. This is an indication of changes in the Trp microenvironment that promotes the quenching of the fluorescence emission of BSA by LT4.

**FIGURE 3.**
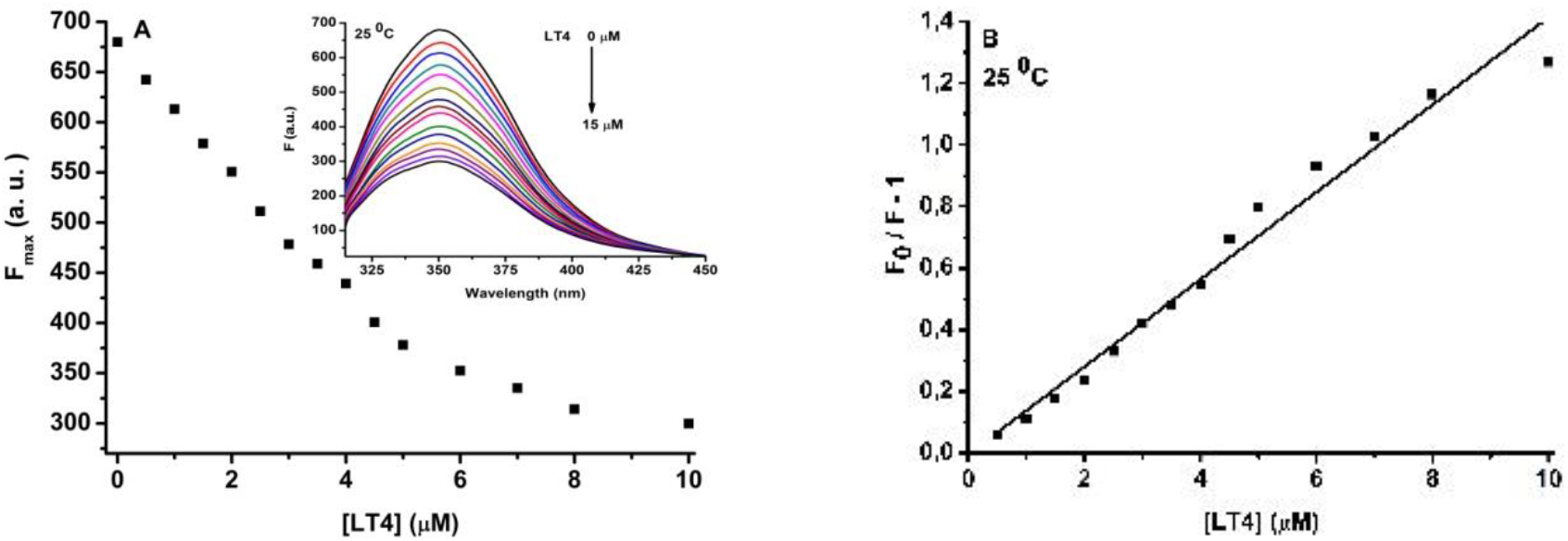
(A) The saturation curve of the BSA (3 μM) molecule by the addition of 0 - 10 μM LT4. Insets: the spectra with gradual titration of LT4 ligand in the BSA solution at 25 °C. Excitation wavelength was 295 nm; samples were diluted in 100 mM HEPES buffer, pH 7.4. (B) The Stern-Volmer representation of the quenching of BSA emission fluorescence by 0 - 10 μM LT4 at 25 ºC.

The binding mechanism of two biomolecules may be static (through the formation of a complex between the quencher molecule and the fluorophore) or dynamic (by a collision processes) (26). In order to determine the nature of the BSA fluorescence quenching by LT4, experimental the data, collected at 25 °C and 35 °C, were analyzed according to the Stern-Volmer eq. (4):

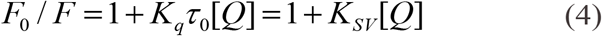

where *F_0_* and *F* are the fluorescence intensity of the protein in the absence and respectively, in the presence of the of quencher, (Q) is the concentration of the quencher, *K_q_* is the bimolecular quenching rate constant, *τ_0_* is the average fluorescence lifetime of the molecule in the absence of the quencher, and *K_SV_* is the Stern-Volmer constant (the quenching constant).

The Stern-Volmer constants were calculated as the ratio of the intercept and the slope for LT4-BSA interaction. The Stern-Volmer plot of the quenching of BSA emission fluorescence by LT4 at 25 °C is shown in Fig. 3 B (similar behaviour was obtained at 35 °C, data not shown). The values of the two Stern-Volmer constants are listed in Table 2. We noticed that the process is not temperature-dependent, thus it is not a dynamic process. We can presume that the quenching mechanism of BSA fluorescence by LT4 is due to the formation of a complex between BSA molecule and LT4 or to a combination with a nonradiative energy transfer process.

**TABLE 2.**
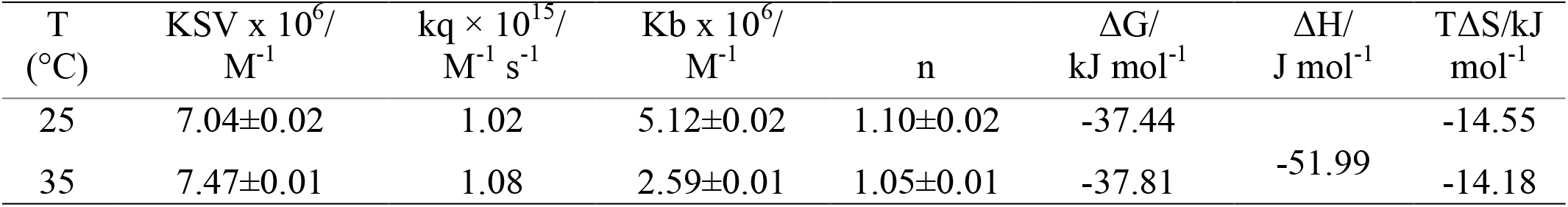
The binding parameters for the interactions of BSA with LT4

Native Trp fluorescence has an average lifetime in apo proteins around τ = 6.9 ns (32). Using this information, the bimolecular constant *k_q_* was calculated (Table 2). The values of this constant are greater than the value for the diffusion limit in aqueous solutions, 1 × 10^10^ M^−1^ s^−1^ (26). Therefore, we can say that LT4 binding with BSA is driven by the complex formation rather than by a collision process.

### 3.4 The strength of the BSA-LT4 binding

The binding constant and the stoichiometry of the BSA-LT4 interaction, studied by intrinsic fluorescence quenching of BSA by LT4, reflect the strength of the interaction. These can be determined using the double logarithmic Scatchard eq. (5) as previously in (23, 24):

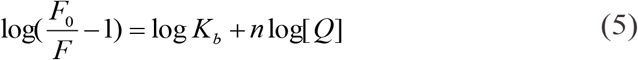

where F_*0*_ and *F* are the fluorescence intensities of BSA in the absence and presence of the LT4, *K_b_* is the binding constant, and *n* is stoichiometry or the number of binding sites of BSA for the ligand.

The plot of log (*F_0_*/*F* - 1) versus log((LT4)) (Fig. 4) allows us to determine the binding constant. We obtained the log *K_b_* from the intercept and *n* from the slope of the plot. The *K_b_* and *n* values obtained at 25 °C and 35 °C are listed in Table 2. One molecule of LT4 binds one molecule of BSA, in accordance with older studies (33), and the binding process is characterized by a moderate interaction (34).

**FIGURE 4.**
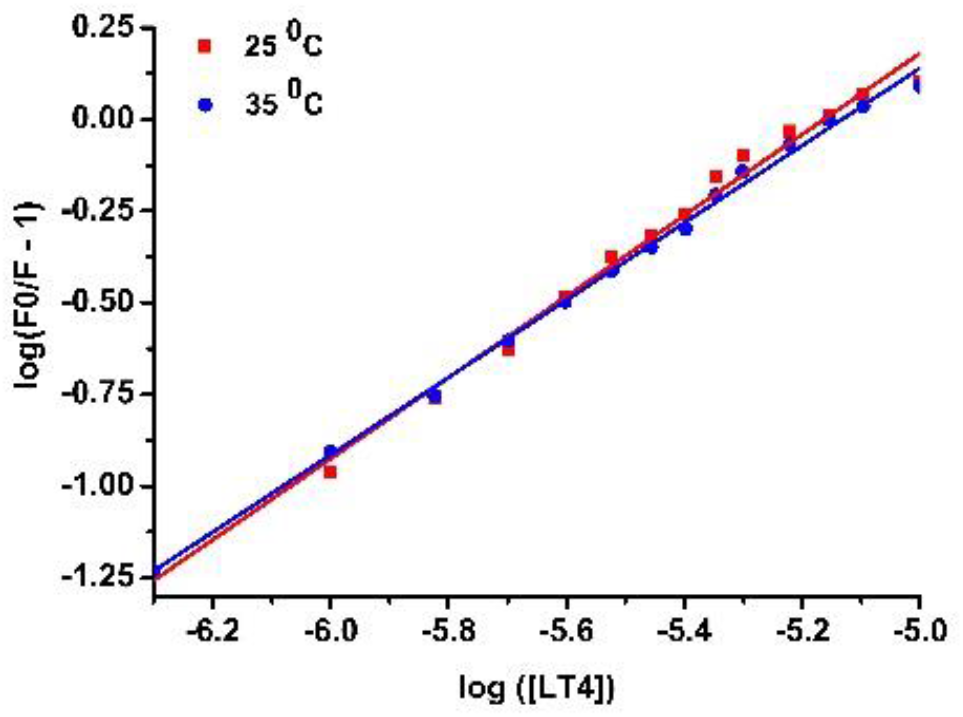
The Scatchard plot of the BSA emission fluorescence quenching by LT4 at 25 °C and 35 °C.

The binding of various ligands transported by BSA to the protein site(s) is reversible. In the case of LT4 binding to the BSA site, the binding (association) constant *K_b_* at 25 °C is 5.12 × 10^6^ M^−1^, and at 35 °C is 2.59 × 10^6^ M^−1^ (Table 2). Therefore, the macromolecular dissociation constant, *K_d_*, of the complex will be 1.95 × 10^−5^ M, at 25 °C and 3.86 × 10^−5^ M at 35 °C.

### 3.5 Thermodynamic parameters and nature of the binding forces between LT4 and BSA

A combination of several forces may be involved in the interaction between a protein and its ligand: hydrogen bonds, electrostatic, hydrophilic/hydrophobic and van der Waals forces. The sign and magnitude of the thermodynamic parameters of the binding are correlated with particular kinds of interactions that may occur in protein association processes (35). Thus, in order to characterize the driving forces for the binding of BSA and LT4, the thermodynamic parameters of the BSA-LT4 interaction were calculated, according of eq. (6) and eq. (7):

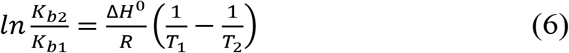

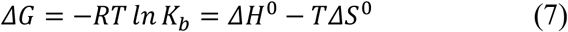

where *K_b_* is the binding constant at respective temperatures; *T* is the absolute temperature; *R* is the gas constant (≅ 8.314 J K^−1^ mol^−1^). *ΔH^0^* and *ΔS^0^* are the variations of enthalpy, respectively entropy in standard conditions.

The binding parameters for the interactions of BSA with LT4 are listed in Table 2. These values are in agreement with previous experiments (36). The interaction of LT4 with BSA is a spontaneous process (*ΔG* < 0). We found that |*ΔH*| > *TΔS* thus the binding of LT4 to BSA is mainly enthalpically driven. Because *ΔH* < 0 and *ΔS* < 0, one can deduce that the hydrogen bonding and van der Waals forces are the main driving forces of the binding process, as other publications suggest (33, 36).

### 3.6 Fluorescence resonance energy transfer (FRET) from BSA to LT4

An energy transfer between two molecules can occur when the fluorescence emission spectrum of the donor overlaps the absorption spectrum of the acceptor, in particular conditions (26). In the case of BSA (3 μM) - the donor and LT4 (3 μM) - the acceptor, the overlap of the two spectra in the range, 310 nm - 500 nm is presented (Fig 5).

**FIGURE 5.**
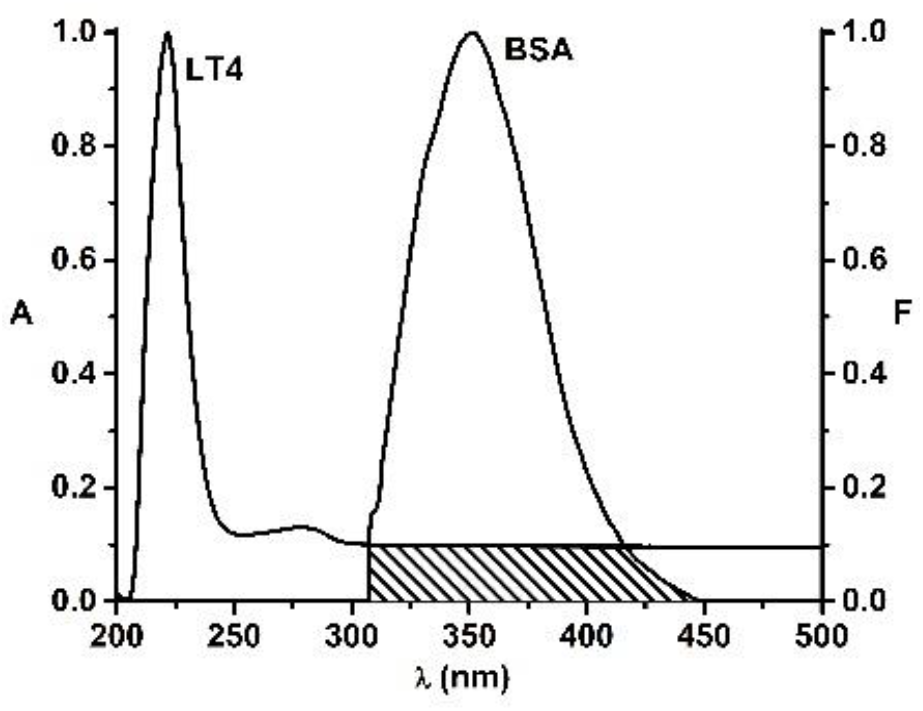
The overlap of the fluorescence emission spectrum of BSA and the absorption spectrum of LT4 ((BSA):(LT4) = 1:1).

We determined the overlap integral of the fluorescence emission spectrum of the donor with the absorption spectrum of the acceptor, *J* = 1.41 × 10^14^ M^−1^ cm^−1^ nm^4^ and we used *k*^2^ = 2/3, *n* = 1.336, and *Q*_D_ = 0.118 (23) in order to determine *R_0_* = 2.59 nm, using the Eq. (2).

The efficiency of energy transfer, *E*, was found to be *E* = 0.55, and the distance between BSA and LT4 molecules was *r* = 2.48 nm. These values confirm the quenching results, which showed that the binding mechanism between BSA and LT4 is static.

### 3.7 The Binding of LT4 to BSA as measured by Surface Plasmon Resonance

Surface plasmon resonance (SPR) studies were performed in order to determine the binding affinity of LT4 towards BSA. SPR enables the study of label-free biomolecular interaction processes, since they lead to small changes in the refractive index in the vicinity of a metal surface (37). Surface plasmons are electromagnetic waves, which are generated due to the differences between the dielectric constant (ε) of each medium at the water/metal interface. The dielectric constant of the metal (ε_1_) must be negative, and its magnitude greater than that of the dielectric constant (ε_2_) of the solution (38). Herewith, in this study, sensitive, quantitative and qualitative measurements were performed in order to monitor both the interaction between confined BSA on SPR chips surface and LT4, and to optimize the immobilization of BSA molecules at chips gold surface. Angular scan measurements (AS-SPR) permit to monitor changes of the angle shift and intensity variation, but also changes of the SPR angle in time (sensograms). All measurements were performed in a continuous flow of 0.1 M HEPES, pH 7.4, with a flow rate of 30 μL/min, except for the functionalization of the gold surface, by Cys adsorption, when the flow was stopped for 20 min. All used analytes were injected manually, and washing of excess molecules occurred through the continuous buffer flow.

#### 3.7.1. Immobilization of BSA onto SPR chips

BSA is widely used in the study of various pharmaceutical products and therefore, its immobilization on different substrate surfaces is an important step that needs to be carefully monitored. Many parameters affect the adsorption /binding of proteins on different substrates, including the nature of the substrates, such as hydrophilicity or hydrophobicity and surface morphology. The interaction capacity of immobilized BSA with LT4 depends on the BSA immobilization method. It is well known that BSA molecule contains one free SH group (39) and therefore we assume this should adhere to the gold surface of the SPR chip. This approach was successfully tested in literature, where direct BSA adsorption onto gold was compared to its adsorption onto different surfaces (40). Although the authors found that a slightly acidic pH of 5.5 is more favourable for BSA adsorption, the neutral pH value of HEPES was kept in order to simulate physiological conditions. Three different concentration of BSA (duplicate measurements) were directly immobilized onto the gold surface, as shown in Fig 6. For higher concentrations of BSA solutions (1 and 10 mg mL^−1^), one can notice significant differences in the signal intensity after the washing step. The best reproducibility was found for the lowest concentration of 0.5 mg mL^−1^ BSA, corresponding to a concentration of injected BSA solution of 7 μM which is close to the concentration used for free BSA during the fluorescence measurements. The SPR Navi Data Viewer software enables to calculate the mass coverage for all concentrations and the results are presented in Table 3.

**FIGURE 6.**
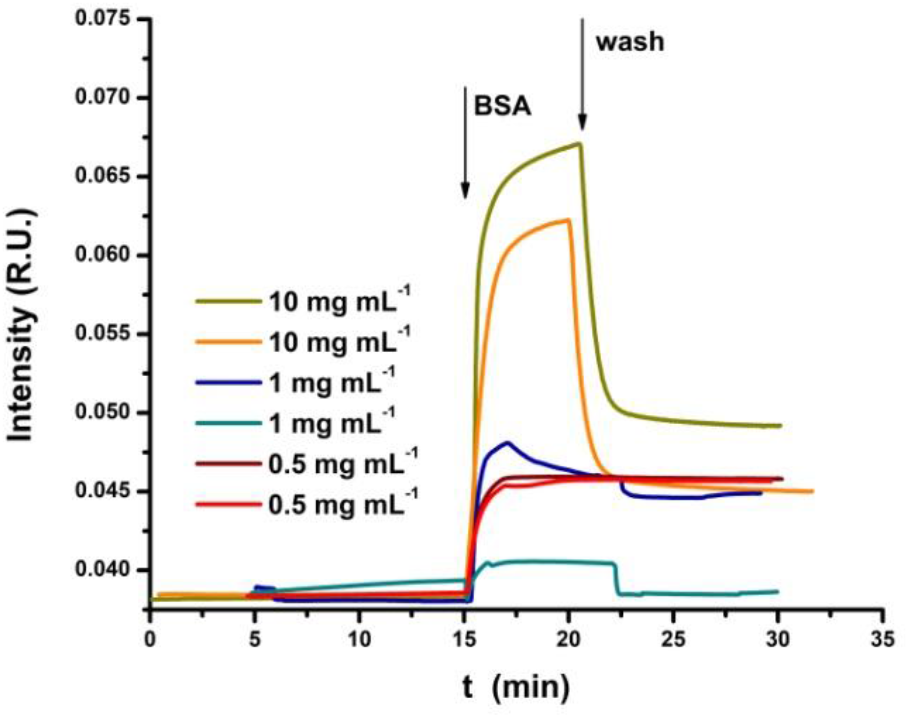
Sensograms for BSA immobilization on SPR chips using different solution concentrations.

**TABLE 3.** Mass coverage estimation for different BSA solution concentrations (n=2) calculated from the sensograms in Fig. 6:

| BSA solution concentrations (mg mL<sup>-1</sup>) | m (ng cm<sup>-2</sup>) |
| --- | --- |
| 10.0 | 35.51 ± 2.39 |
| 1.0 | 6.16 ± 5.08 |
| 0.5 | 10.91 ± 0.29 |

When a solution with 0.5 mg mL^−1^ was used, the adsorption of BSA was reproducible and the washing step does not seem to remove any of the unbound molecules. For the rest of situations, the direct BSA immobilization on gold surface was not reproducible. Herewith, we conclude that direct BSA adsorption is not stable enough on SPR chips to be used for further experiments. Therefore, a low concentration of BSA should be immobilized onto the gold surface using linkers. A few successful examples can be found in literature, most of them making use of carboxymethyl dextran (CMD) modified gold surfaces, activated by NHS/EDC chemistry, followed by BSA covalent amide linking (41, 42). Based on these findings, we chose two approaches to immobilize BSA using the solution of 0.5 mg mL^−1,^ which correspond to an adsorbed mass of 10.91 ng cm^−2^ (165 nM).

First, we used commercially available −NH_2_ modified gold chips (Bionavis, Finland) and the immobilization method is denoted M1. All solutions were injected manually during a continuous buffer flow. In order to activate the amino moieties on the chip, an aqueous 1 % GA solution was used, as shown in the sensograms from Fig 7A. All aqueous solutions were prepared using 0.1 M HEPES buffer, pH 7.4, taking into account that SPR is very sensitive to the changes in solution parameters. Since the measurements were done in a continuous flow, no additional washing steps were required, each injected solution being carried away with the flow. After the activation of −NH_2_ group, the immobilization of BSA followed through amine coupling. In order to block non-specific binding on the chip surface, 1M of ETA was used.

**FIGURE. 7.**
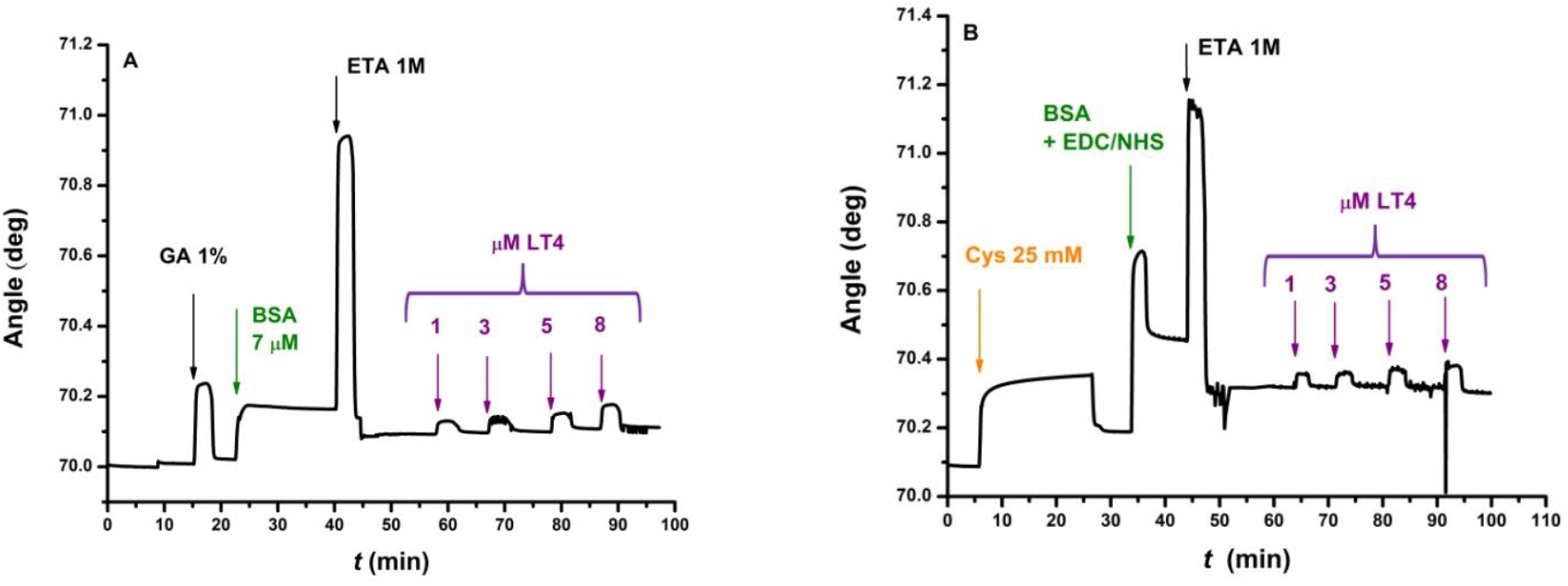
Sensogram of BSA immobilization process on gold chip followed by consecutive LT4 injections with increasing concentrations, using two different chips: (A) Au-NH_2_-GA-BSA and (B) Au-Cys-BSA.

We can clearly see how the unbound BSA molecules will be removed, as well as the inactivation of unoccupied −NH_2_ groups, creating a stable BSA film for further LT4 binding. The second approach to immobilize BSA on the gold surface, denoted M2, is presented in Fig 7B. In this case, the bare gold chip was functionalized with −NH_2_ moieties, based on previous findings (43), using a 25 mM Cys solution in static condition for 20 min. In order to assure the proper linking through Au-S bonds, the buffer flow was stopped.

After Cys adsorption, BSA immobilization followed. BSA was mixed with EDC/NHS solution overnight in the refrigerator in order to activate the carboxyl groups (-COOH) present in the protein (EDC and NHS were mixed in a ratio of 4:1 corresponding to 40 mM EDC and 10 mM NHS). 1 M ETA was used to block non-specific binding on the chip surface.

#### 3.7.2 Binding affinity of LT4 towards BSA

After optimizing the two immobilization methods for BSA on SPR chip surfaces, its interaction with different LT4 concentrations (3 - 20 μM) was monitored (Fig. 7). Each sample of different concentration was injected to the BSA-modified surface, and the variation of the peak minimum angle measured at chemical equilibrium was plotted as a function of LT4 concentration. All measurements were done in triplicate. From a pharmacological and biochemical point of view, the binding of ligands to macromolecules as a function of ligand concentration are well described by the Hill-Langmuir equation which is used to quantify the ligand-receptor interaction (44):

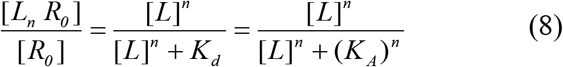

where (*L_n_R*)/(*R_0_*), often denoted θ, represents the fraction of the receptor protein sites - bound by the ligand, (*R_0_*) is the total amount of receptor concentration, (*L*) and (*L_n_R*) represent the free ligand and ligand-bound receptor concentrations, *K_d_* is the *apparent* equilibrium dissociation constant derived from the law of mass action, *K_A_* denotes the microscopic dissociation constant (it is the ligand concentration at half occupation of receptor sites) and *n* (also denoted *n_H_*) is the Hill coefficient representing the number of binding sites (if *n_H_* = 1, *K_A_* equals *K_d_*).

The Hill-Langmuir equation was used to determine the binding affinity of LT4 towards confined-BSA using the two immobilization methods described above. The concentration dependent calibration plot is shown in Fig. 8A, where the SPR angle increases with LT4 concentration (only for the M2 chip with similar behaviour for M1 chip). The linearization of this equation is obtained by rearranging eq. (8) to:

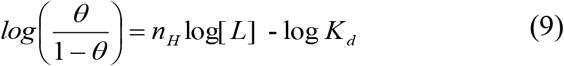

where plotting *log(*θ/(1-θ)) versus log *(L)*, the Hill plot, is obtained (Fig 8B). The slope represents the Hill coefficient, *n_H_*, and the intercept, *log (K_d_),* provides the *apparent* dissociation constant *K_d_* (45).

**FIGURE 8.**
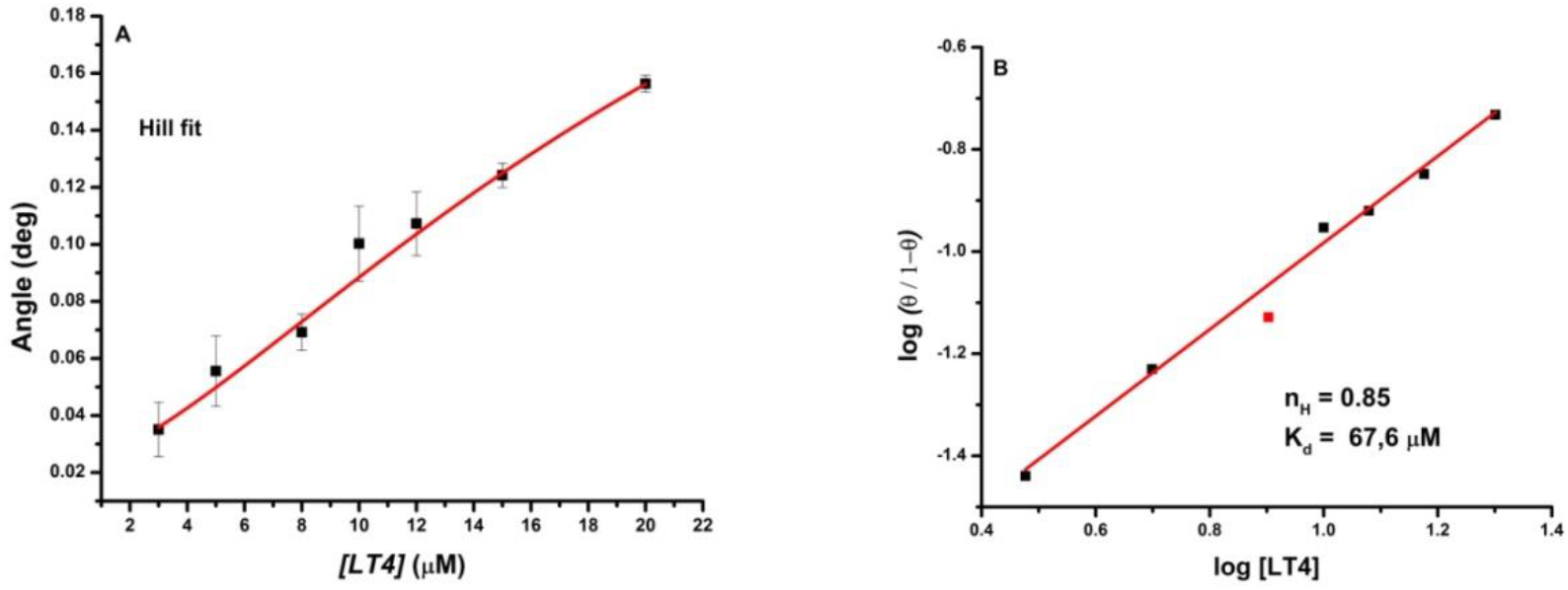
(A) Calibration plot for Au-Cys-BSA chip in the presence of increasing LT4 concentrations. (B) Corresponding Hill plot.

All parameters were calculated for both BSA immobilization methods and their values are presented in Table 4. The *apparent* dissociation constant, *K_d_*, corresponds for BSA-LT4 complex confined at SPR chip surface and the Hill coefficient gives an insight of the binding cooperativity. Low *K_d_* values suggest a strong binding affinity, and in our case, the M2 immobilization method suggests a more suitable orientation of the BSA molecules towards LT4 binding. However, if compared to other *K_d_* values in literature (46, 47), where the dissociation constant range is of the order of nM, respectively pM, one can conclude that the binding affinity between confined-BSA and LT4 is weak. This is also in accordance with the findings made through fluorescence measurements, which indicate that the BSA-LT4 binding process is characterized by a moderate interaction. Thus, the *apparent* constants of dissociation obtained by SPR are higher than the dissociation constant obtained in the fluorescence experiment (*K_d_* = 0.195 μM). This difference may be due to BSA immobilization on SPR chips, as opposed to fluorescence, where the protein site has a different exposure towards LT4, exposure influenced also by the interaction with the solvent.

**TABLE 4.** Binding affinity parameters determined for both BSA immobilization methods

| Methods | $K_d$ ($\mu\text{M}$) | $n_H$ | $r^2$ |
| --- | --- | --- | --- |
| M1 | 74.1 | 1.05 | 0.966 |
| M2 | 67.6 | 0.85 | 0.995 |

By analyzing the value of the Hill coefficient, the cooperativity of the binding can be determined. Since BSA presents more than one binding sites, the Hill coefficient should be regarded as an “interaction” coefficient describing the cooperativity among multiple binding sites (48). For M1, *n_H_* is very close to one, suggesting a noncooperative binding, where the *apparent* dissociation constant *K_d_* can be regarded as *K_A_*, and the binding process can be modelled after the Michaelis-Menten kinetics. The BSA-LT4 binding is similar to a substrate-enzyme association which is reversible, specific and saturable (43). For the second immobilization method, *n_H_* is lower than 1, suggesting a negatively cooperative binding. Thus, we can say that BSA exhibits competitive binding characteristics, which is in accordance with the findings throughout this work.

Summarizing, M2 is most suitable for BSA immobilization and further proof in this perspective is provided by the SPR angular scan (AS-SPR) measurements from Fig 9. Here, the resonance of the surface plasmons is observed as a shadow in the reflected light at the SPR angle, which depends on the mass of the dielectric layer at the gold surface. When biomolecules bind to the gold surface, the resonance angle will shift towards higher values, as can be seen in Table 5. Changes in intensity suggest the presence of light adsorbing molecules or molecule complexes: the more light is adsorbed, the more light adsorbing molecules are present on the chip surface.

**FIGURE 9.**
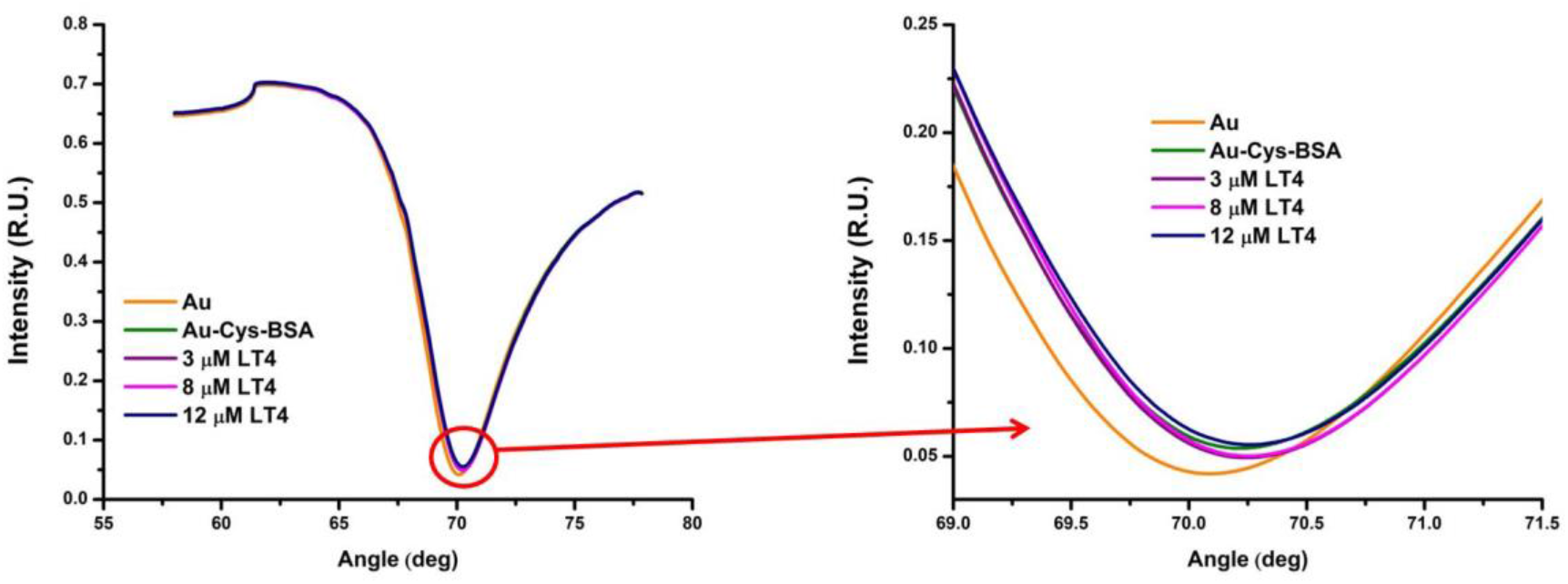
AS-SPR monitoring of biomolecular interactions onto Au-Cys-BSA chip followed by titration of LT4.

**TABLE 5.**
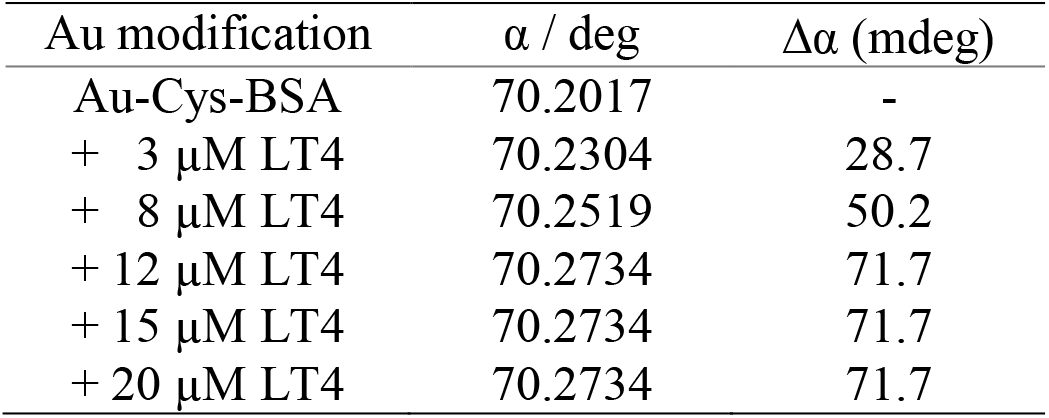
SPR angle shift (Δα) upon LT4 titration using Au-Cys-BSA chip

| Au modification | $\alpha$ / deg | $\Delta\alpha$ (mdeg) |
| --- | --- | --- |
| Au-Cys-BSA | 70.2017 | - |
| + 3 $\mu\text{M}$ LT4 | 70.2304 | 28.7 |
| + 8 $\mu\text{M}$ LT4 | 70.2519 | 50.2 |
| + 12 $\mu\text{M}$ LT4 | 70.2734 | 71.7 |
| + 15 $\mu\text{M}$ LT4 | 70.2734 | 71.7 |
| + 20 $\mu\text{M}$ LT4 | 70.2734 | 71.7 |

With the addition of increasing LT4 concentrations, the SPR angle shifts towards higher degrees as compared with Au-Cys-BSA value. For higher concentrations than 12 μM LT4, the angle remains constant. Thus, the angle variation, Δα, reached a constant value. In the case of reflected intensity, during the chip surface modification, the reflected intensity decreases, proving the BSA immobilization (light absorption). Thus, the changes in angle shifts and reflected intensity provide information on the BSA immobilization onto chip surface and confined-BSA-LT4 complex formation.

### 3.8 Spectroscopic biosensors for LT4 detection

By analyzing the data from the above sections we can say that BSA-LT4 interaction can be used to develop biosensors for LT4 detection. The Stern-Volmer representation of the quenching of BSA fluorescence emission by 0 - 10 μM LT4 at 25 °C, a sensitivity of *S* = 59.03 ± 1.4 a.u. μM^−1^ with a limit of detection, LOD = 0.23 μM, were obtained. The calibration plot of the fluorescence biosensors for LT4 is shown in Fig. 10.

**FIGURE 10.**
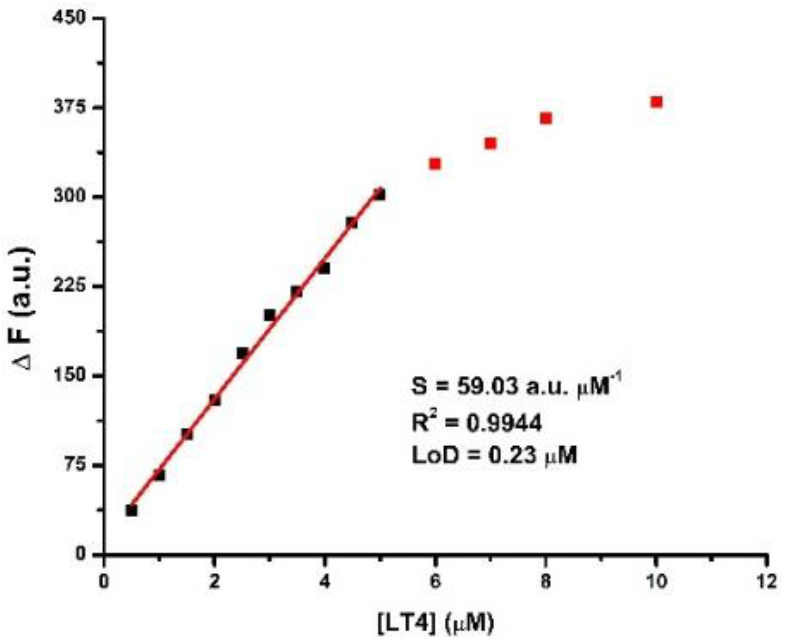
Calibration plot for LT4 titration.

In the cases of confined-BSA, as possible SPR biosensor for LT4 detection, two values can be used. One way can be provided by plotting the estimated mass coverage of LT4 at Au-Cys-BSA SPR chip (Δm) as a function of LT4 concentration (data conversion from sensograms in Fig. 7B). A sensitivity, S = 7.01 ± 0.3 ng cm^−2^ μM^−1^ with an LOD of 0.37 μM has been obtained. A second modality is given by the plot of the SPR angle shift (Δα) as a function of the LT4 concentration (data from Fig. 9), where a sensitivity, *S* = 5.93 ± 0.31 mdeg μM^−1^ and a limit of detection LOD = 1.38 μM was calculated. In both cases, the calibration plot is similar with that from Fig. 10 (data not shown). For all situation, LODs have been calculated as (3·x SD/S), where SD stands for standard deviation of the intercept. All measurements were done in triplicate.

We can see that by fluorescence quenching of free BSA (monitoring fluorescence intensity variation) and SPR of confined-BSA (sensogram monitoring variation of bound LT4 mass) both techniques can be used for biosensing of LT4 in the nanomolar range.

## Conclusion

Binding of the hydrophilic compound LT4 to BSA can significantly alter the pharmacological properties of the compound, and therefore, studying their interaction could be a difficult issue, depending on the methods used for testing. The binding mechanism and affinity of the interaction between LT4 and free BSA were investigated in solution by FT-IR, fluorescence, and FRET. Confined-BSA on Au chip and its interaction with LT4 was studied using SPR.

Using the UV-Vis technique the value of 7.4 was found to be the optimal pH for LT4 in HEPES buffer.

Fluorescence experiments showed that LT4 interacts with BSA mainly by a static mechanism, and affects the protein conformation at the binding site. Binding is a spontaneous process, induced by van der Waals forces and formation of hydrogen bonds (*K_b_* = 5.12 μM^−1^, respectively *K_d_* = 0.195 μM).

Using SPR, BSA was confined to Au surface by two immobilization methods. The binding affinity of LT4 towards confined-BSA activated by NHS/EDC chemistry, followed by BSA covalent amide linking was found to have better characteristics determined by Hill-Langmuir equation, which suggest an *apparent K_d_* = 67.6 μM and a negatively cooperative binding (*n_H_*<1). The difference between this *apparent* dissociation constant value and the one obtained using fluorescence may be due to BSA immobilization on SPR chips, and indicates that the diffusion limitations inside the BSA layer immobilized on the chip surface play an important role. The confinement influences both protein conformation and the binding site exposure towards LT4.

Both fluorescence quenching of free BSA and SPR of confined-BSA can be used as biosensors for LT4 detection in the nanomolar range. As SPR was applied for the first time to study LT4 – protein interaction, further studies will lead to the development of sensitive spectroscopic methods with clinical and pharmacological applications, both for the characterization of LT4 protein-drug interaction and LT4 detection.

## Author Contributions

Nicoleta Sandu: Investigation, Resources, Writing - original draft. Claudia G. Chilom: Conceptualization, Validation, Formal analysis, Investigation, Resources, Writing - original draft. Melinda David: Conceptualization, Methodology, Validation, Formal analysis, Investigation, Writing - original draft. Monica Florescu: Conceptualization, Methodology, Validation, Formal analysis, Investigation, Resources, Writing - review & editing.

## Acknowledgements

This work was supported by a grant of the Romanian Ministry of Research and Innovation, CCCDI - UEFISCDI, project number PN-III-P1-1.2-PCCDI-2017-0062, contract no. 58, within PNCDI III and the structural funds project PRO-DD (POS-CCE, O.2.2.1., ID 123, SMIS 2637, No 11/2009) for providing some of the infrastructure used in this work. The authors are grateful to Dr. George Stan and Dr. Adrian Enache (National Institute of Materials Physics, Laboratory of Multifuntional Materials and Structures, Magurele, Romania) for FT-IR spectra and are very much indebted to Prof. Dr. Aurel Popescu, for very helpful suggestions and permanent encouragement.

